# The evolution of the cytochrome *c*_6_ family of photosynthetic electron transfer proteins

**DOI:** 10.1101/2021.02.16.431308

**Authors:** Barnaby Slater, Darius Kosmützky, R. Ellen R. Nisbet, Christopher J. Howe

## Abstract

During photosynthesis, electrons are transferred between the cytochrome *b*_6_*f* complex and photosystem I. This is carried out by the protein plastocyanin in plant chloroplasts. In contrast, electron transfer can be carried out by either plastocyanin or cytochrome *c*_6_ in many cyanobacteria and eukaryotic algal species. There are three further cytochrome *c*_6_ homologues: cytochrome *c*_6A_ in plants and green algae, and cytochromes *c*_6B_ and *c*_6C_ in cyanobacteria. The function of these proteins is unknown. Here, we present a comprehensive analysis of the evolutionary relationship between the members of the cytochrome *c*_6_ family in photosynthetic organisms. Our phylogenetic analyses show that cytochrome *c*_6B_ and cytochrome *c*_6C_ are likely to be orthologues that arose from a duplication of cytochrome *c*_6_, but that there is no evidence for separate origins for cytochrome *c*_6B_ and *c*_6C_. We therefore propose re-naming cytochrome *c*_6C_ as cytochrome *c*_6B_. We show that cytochrome *c*_6A_ is likely to have arisen from cytochrome *c*_6B_ rather than by an independent duplication of cytochrome *c*_6_, and present evidence for an independent origin of a protein with some of the features of cytochrome *c*_6A_ in peridinin dinoflagellates. We conclude with a new comprehensive model of the evolution of the cytochrome *c*_6_ family which is an integral part of understanding the function of the enigmatic cytochrome *c*_6_ homologues.

## Introduction

### The cytochrome *c*_6_ family of proteins

Photosynthesis is one of the most important processes in the natural world and has played a vital role in shaping the planet and its atmosphere. One essential feature of oxygenic photosynthesis is the photosynthetic electron transfer chain (PETC), where the oxidation of water to generate reducing equivalents and chemical energy as ATP is driven through light energy absorption. In the plant PETC, electrons can be transferred between the cytochrome *b*_6_*f* complex and photosystem I by the copper-containing protein plastocyanin (Gross, 1993). Many cyanobacteria and eukaryotic algae have an alternative electron transfer protein to plastocyanin, the haemoprotein cytochrome *c*_6,_ which is used when copper is not readily available (Wood, 1978). It is believed that cytochrome *c*_6_ is a more ancient protein than plastocyanin, with the latter evolving after increasing atmospheric oxygen concentrations led to a decrease in the ready availability of iron in the environment (De la Rosa *et al*., 2002). Higher plants were believed to have lost cytochrome *c*_6_, retaining only plastocyanin (Kerfeld and Krogmann, 1998).

However, in 2002 a homologue of cytochrome *c*_6_ was found in higher plants (Gupta, He and Luan, 2002; Wastl, Bendall and Howe, 2002). This protein was subsequently named cytochrome *c*_6A_ (Wastl *et al*., 2004; figure 1). The sequence of cytochrome *c*_6A_ was found to be highly similar to that of *c*_6_, with a major difference that cytochrome *c*_6A_ contains a 12-amino acid insertion in a loop region of the protein. This insertion has been named the loop insertion peptide (LIP) (Howe *et al*., 2006), and contains two cysteines that form a disulphide bridge (Marcaida *et al*., 2006). Further homologues of cytochrome *c*_6A_ (in addition to the conventional cytochrome *c*_6_) were then discovered in cyanobacteria, and named cytochromes *c*_6B_ and *c*_6C_ (Nomura, 2001; Bialek *et al*., 2008; figure 1). These cytochromes were split into B and C homologues based on a phylogenetic analysis, which showed that cytochrome *c*_6B_ shared a more recent common ancestry with cytochrome *c*_6A_, and cytochrome *c*_6C_ shared a more recent common ancestor with cytochrome *c*_6_ (Bialek *et al*., 2008).

**Figure 1.**
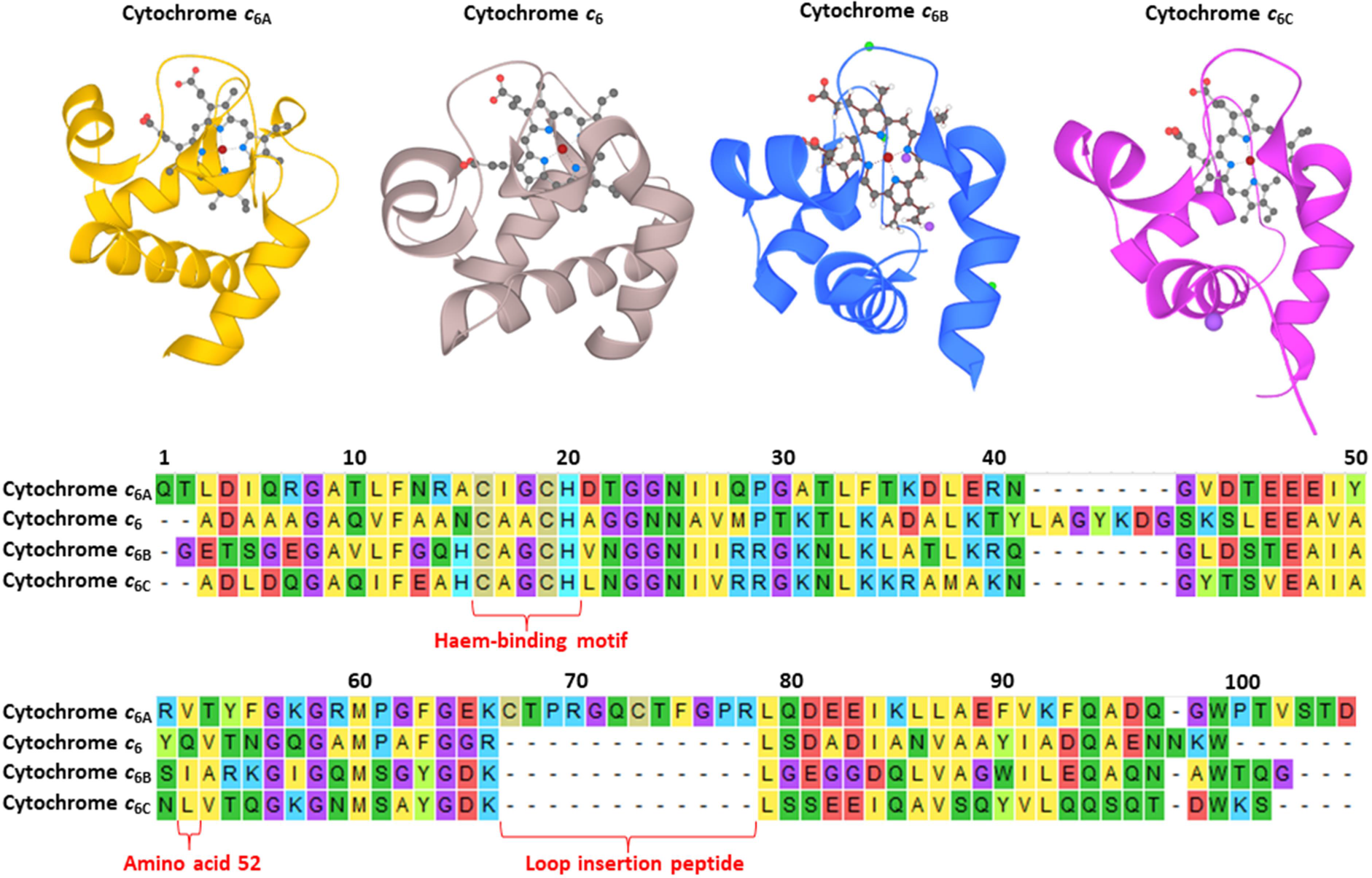
Top, X-ray crystal structures of cytochromes *c*_6A_ (yellow), *c*_6_ (beige), *c*_6B_ (blue) and *c*_6C_ (magenta). Secondary structure for each protein is shown in ribbon form, and the haem prosthetic groups shown in ball and stick (carbon-black, oxygen-red, nitrogen-blue, iron-deep red and sodium-purple). Bottom, protein sequence alignment of cytochromes *c*_6A_, *c*_6_, *c*_6B_ and *c*_6C_ from Arabidopsis thaliana (accession Q93VA3.1), Synechococcus sp. PCC 7002 (accession O30881.1), Synechococcus WH8103 (CRY92441.1) and Synechococcus sp. PCC 7002 (accession AAN03578.1) respectively. The sequences have their putative signal peptides excluded. Amino acids are coloured with yellow-hydrophobic residues, green-polar residues, beige-cysteines, blue-positively charged residues and red-negatively charged residues, and the haem-binding motif (CXXCH), the LIP, and amino acid 52 are indicated below the alignment. Figure uses crystallography and sequence data from Marcaida *et al*., 2006; Bialek *et al*., 2009; Bialek and Jaskolski, 2014; Zatwarnicki *et al*., 2014 (Q93VA3, Q7U624 and Q8KX15 on Uniprot respectively).

Cytochromes *c*_6A_, *c*_6B_ and *c*_6C_ have a redox midpoint potential around 200 mV lower than cytochrome *c*_6_, suggesting that cytochromes *c*_6A_, *c*_6B_ and *c*_6C_ are unable to oxidise cytochrome *f* and have a different function from cytochrome *c*_6_ (Molina-Heredia *et al*., 2003; Bialek *et al*., 2008, 2014). This suggestion of a difference in function was supported by studies on the reaction between cytochrome *c*_6A_ and Photosystem I *in vitro* and the demonstration that plastocyanin is essential in plants (Molina-Heredia *et al*., 2003; Weigel *et al*., 2003). The difference in redox midpoint potential between cytochrome *c*_6A_ and cytochrome *c*_6_ is proposed to be largely due to a single amino acid residue, found at position 52 in *Arabidopsis thaliana* cytochrome *c*_6A_ (Marcaida *et al*., 2006; figure 1). In the low redox midpoint potential cytochrome *c*_6_-like proteins this residue is hydrophobic (leucine, isoleucine or valine), with cytochrome *c*_6_ having a conserved glutamine in the same position.

Substituting the *A. thaliana* cytochrome *c*_6A_ valine 52 with a glutamine has been shown to increase the redox midpoint potential of the protein by around 100 mV (Worrall *et al*., 2007). The function of these low redox midpoint cytochrome *c*_6_-like proteins is currently unclear, though a role in alternative pathways in electron transfer has been proposed (Howe *et al*., 2016).

### The current model of cytochrome *c*_6_ family ancestry

The current hypothesis for the evolution of the cytochrome *c*_*6*_ family in photosynthetic organisms has been outlined by Howe *et al*., 2016 (Figure 2). The model suggested that duplication(s) of cytochrome *c*_6_ in an ancestral cyanobacterium led to the genesis of the low redox midpoint potential cytochromes grouped under the umbrella term cytochrome *c*_6ABC_. Phylogenetic analysis by Bialek *et al*., 2008 suggested at least two duplications had occurred in cyanobacteria, resulting in cytochromes *c*_6B_ and *c*_6C._ Following primary endosymbiosis, cytochrome *c*_6_ was lost in the green plant lineage leaving only a low redox midpoint potential sequence, cytochrome *c*_6A_. Secondary endosymbiosis involving the green lineage, (e.g. as seen for *Euglena)*, was believed to have failed to transfer the low redox midpoint potential cytochrome *c*_6_. In contrast, cytochrome *c*_6ABC_ was probably lost in the red algal and glaucophyte lineages (which contain primary plastids) sometime after the origin of the haptophytes (containing a secondary plastid, (Yoon *et al*., 2002)), which retain both cytochrome *c*_6_ and *c*_6ABC_.

**Figure 2.**
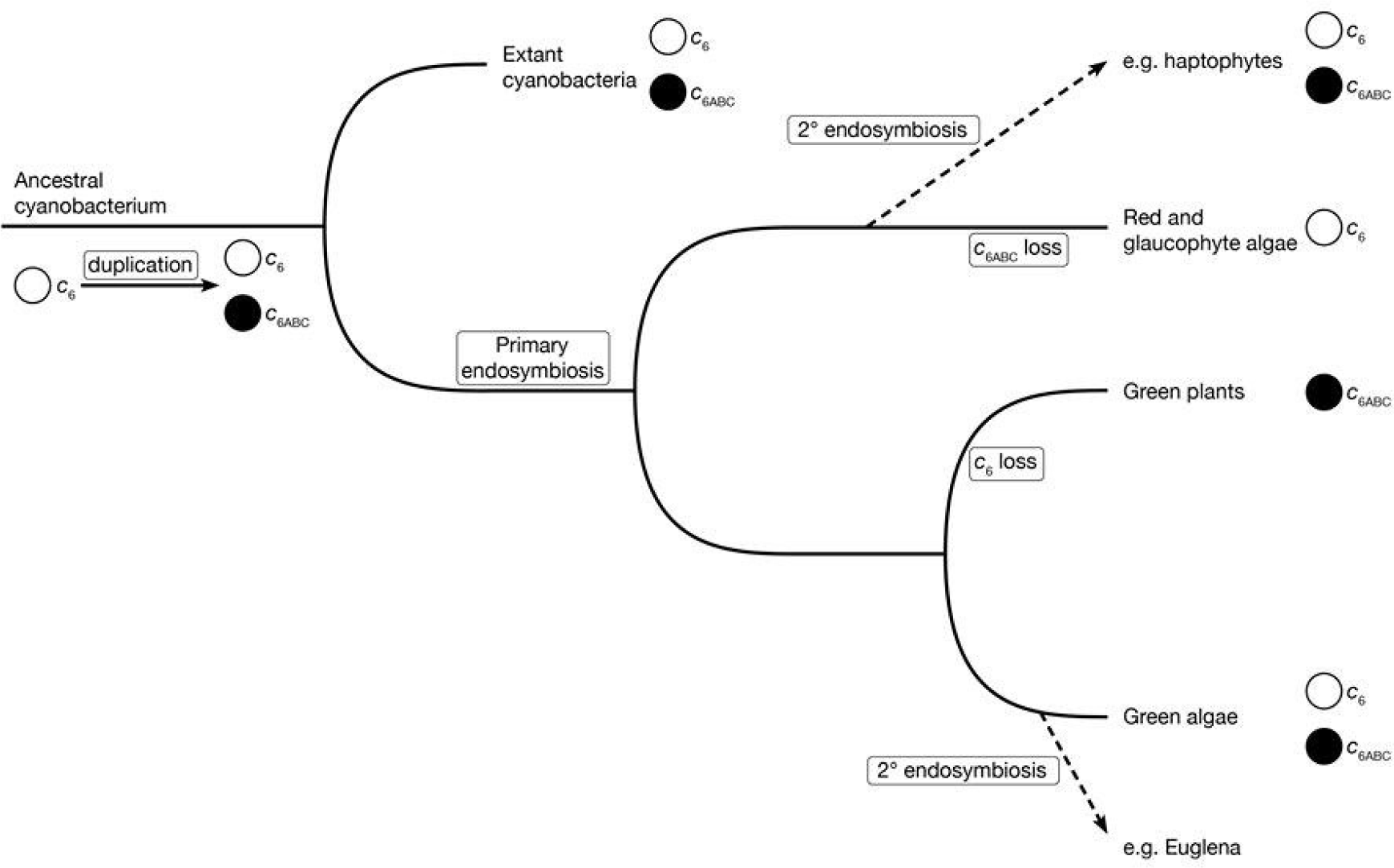
Current model of cytochrome *c*_6_ family evolution in photosynthetic organisms. Adapted from Howe *et al*., 2016.

### Aims of the study

With the availability of more sequence data, this study expanded the search for cytochrome *c*_6_ family sequences in a wider range of photosynthetic taxa, both prokaryotic and eukaryotic. We particularly wished to identify whether cyanobacterial cytochrome *c*_6BC_ proteins were derived from cytochrome *c*_6_ or *vice versa*, what the evolutionary relationship is between cytochrome *c*_6B_ and cytochrome *c*_6C_, and how widely distributed the cytochrome *c*_6ABC_ family is among eukaryotes.

## Results

### Mapping cytochromes *c*_6_, c_6B_ and c_6C_ on an established cyanobacterial species tree shows that *c*_6B_ and *c*_6C_ are orthologues that arose from a single *c*_6_ gene duplication event

To examine the distribution of cytochromes *c*_6_, *c*_6B_ and *c*_6C_ across cyanobacteria, the presence or absence of *c*_6_ family cytochrome sequences was mapped onto a phylogenetic tree of cyanobacterial species inferred from a concatemer of conserved sequences (Walter *et al*., 2017; figure 3). Putative cytochrome *c*_6B/C_ sequences were found by database searching and defined as cytochrome *c*_6B/C_ if they had both an appropriately located haem binding motif (CXXCH; Barker and Ferguson, 1999) and a valine, leucine or isoleucine rather than glutamine at the equivalent of position 52 in *A. thaliana*. (The presence of valine at this position was linked to a lower redox midpoint potential relative to cytochrome *c*_6_ (Worrall *et al*., 2007; Bialek *et al*., 2008)). The sequences identified by Bialek *et al*. (2008) as cytochrome *c*_6B_ were found exclusively in one clade containing cyanobacteria of the genera *Prochlorococcus* or *Parasynechococcus*. The sequences identified by Bialek *et al*. (2008) as cytochrome *c*_6C_ were widespread across the cyanobacterial species tree, but not found in the genera *Prochlorococcus* or *Parasynechococcus*. In addition, no species in the tree contained more than one cytochrome *c*_6B/C_-like sequence. These observations suggest that the prior separation observed between the two sequences could be accounted for by taxon sampling, without needing to propose them as two separate families.

**Figure 3.**
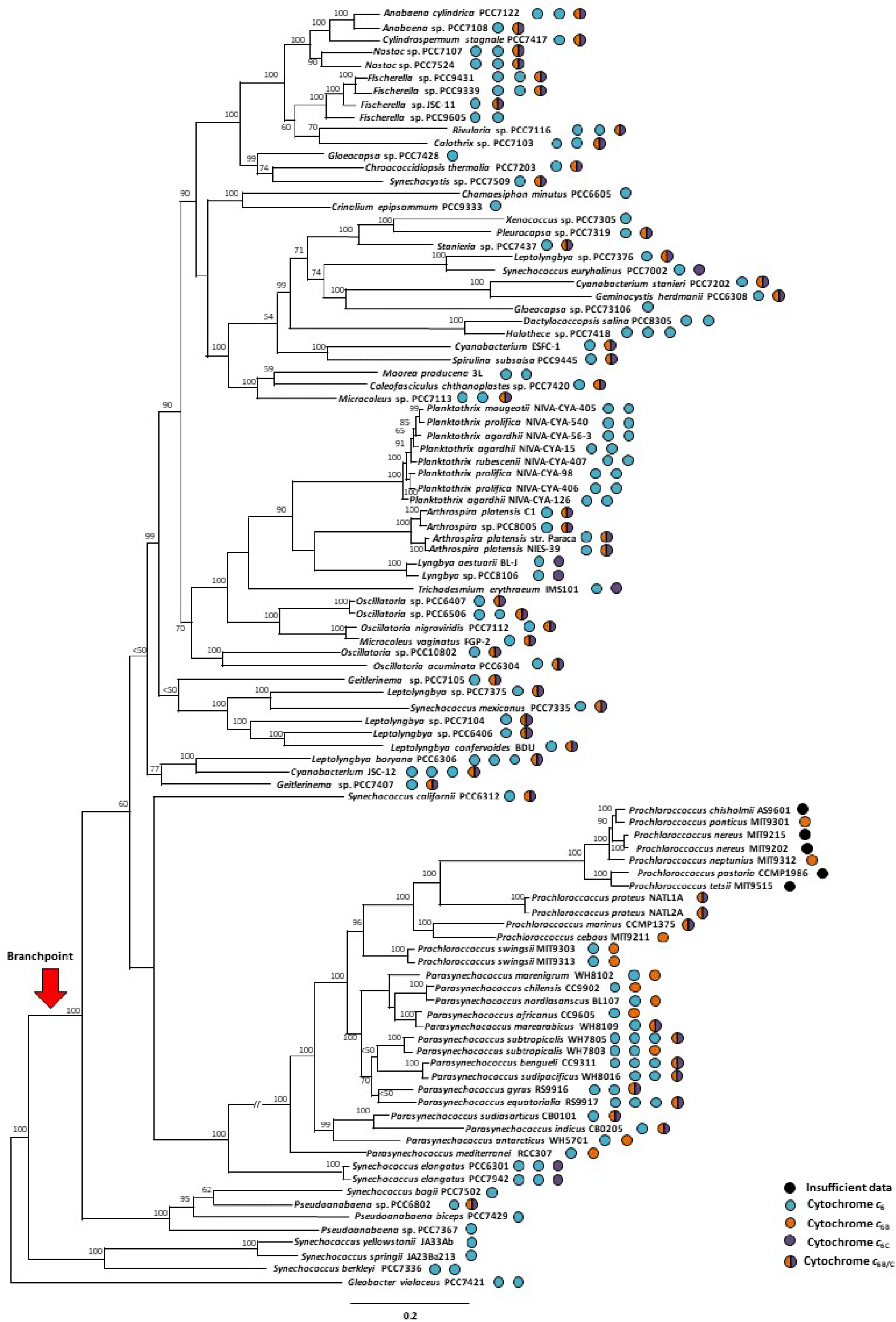
Cyanobacterial species tree with presence of cytochrome *c*_6_, *c*_6B_ and *c*_6C_ mapped onto it (coloured blue, orange and purple respectively). Circles with both orange and purple semi-circles contain a putative low redox midpoint potential cytochrome *c*_6_ which was not included in Bialek *et al*., 2008. Proteins represented by orange or purple full circles were described as *c*_6B_ or *c*_6C_ respectively by Bialek *at al*., 2008. Tree is recreated from Walter *et al*., 2017. The potential branch where neofunctionalisation led to the evolution of cytochrome *c*_6B_ is indicated with a red arrow. Scale bar represents branch length. Bootstrap values calculated using 1000 iterations are marked by the relevant branch point. Accessions of the sequences used for this figure can be found in supplementary table 1.

The cyanobacterial tree shows a branch point (labelled with a red arrow) separating taxa which have only cytochrome *c*_6_ from those which also have a cytochrome *c*_6B_ or *c*_6C_ sequence. This branch point separates the basally-diverging *Gloeobacter violaceus* PCC7421 (Criscuolo and Gribaldo, 2011; Mareš *et al*., 2013) and a few other taxa from the rest. This distribution suggests that cytochrome *c*_6_ appeared first, and that cytochromes *c*_6B_ and *c*_6C_ may have arisen through duplication and neofunctionalisation of cytochrome *c*_6_.

### A phylogenetic tree using a wider taxon selection suggests a single origin for cytochromes *c*_6B_ and *c*_6C_

To see whether any evidence for separate origins of cytochromes *c*_*6B*_ and *c*_*6C*_ within the cytochrome *c*_6_ radiation could be seen with a wider range of sequences, and using a phylogenetic tree constructed on the basis of cytochrome *c*_6_ family proteins, a tree was inferred from an alignment of cytochromes *c*_6_, *c*_6B_ and *c*_6C_ sequences covering all the organisms used in two independent phylogenetic analyses of the cyanobacterial lineage (Schirrmeister et al., 2015; Walter et al., 2017). A condensed tree is shown in figure 4. The cytochromes predicted to have a low redox midpoint potential, including those assigned as cytochromes *c*_6B_ and *c*_6C_ previously, all grouped to the exclusion of the predicted cytochrome *c*_6_ sequences (bootstrap value of 84%), and maintained a similar general topology to that of the cyanobacterial tree of figure 3. This distribution showed cytochrome *c*_6B_ as a clade derived from within the cytochrome *c*_6C_ clade, as with figure 3. Once again, there was no evidence of both a cytochrome *c*_6B_ and *c*_6C_ within the same organism. (*Crocosphaera watsonii* has been shown to have two low redox midpoint potential cytochrome *c*_6_ sequences, but both were assigned as cytochrome *c*_6C_ in prior studies (Bialek *et al*., 2008)). These observations suggest a single origin for the cytochrome *c*_6BC_ family, and that they are orthologues rather than paralogues. It is worth noting that the bootstrap values in this tree were considerably lower than those in the tree inferred by Walter *et al*., 2017, which is most likely due to the short sequence length of cytochrome *c*_6_ family peptides.

**Figure 4.**
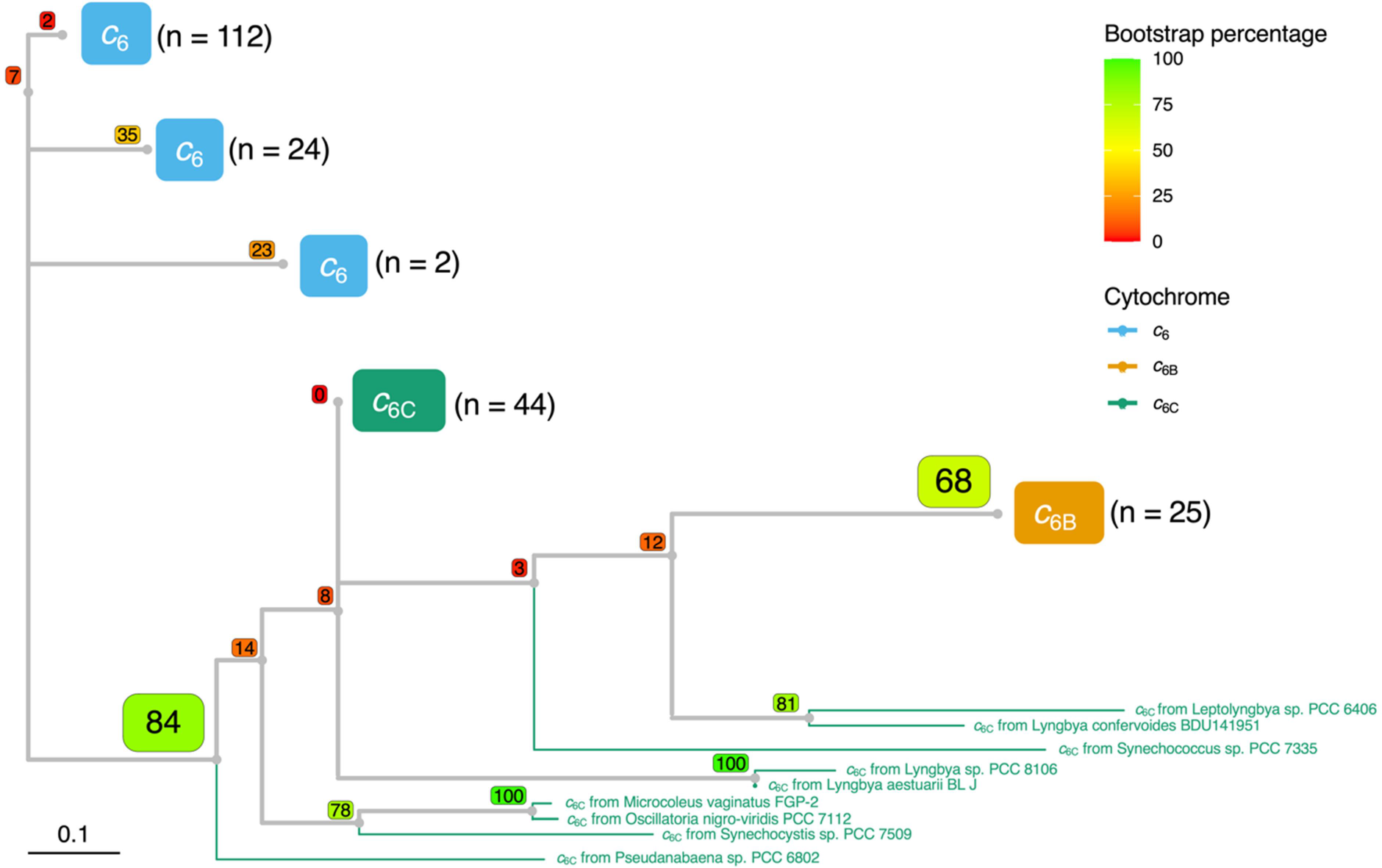
Condensed phylogenetic tree inferred from an alignment of cytochrome *c*_6_, *c*_6B_ and *c*_6C_ peptide sequences from cyanobacterial species (coloured blue, orange and green respectively). Alignments were performed using Muscle algorithm and can be found in the supplementary information along with accessions for each sequence used. The tree was built using maximum likelihood inference using a WAG model with Gamma distribution and invariant sites (WAG+G+I). Bootstrap values for each branch point, using 100 iterations, are shown in coloured boxes. The n value next to each group represents the number of sequences found within each clade. The full tree is shown in supplementary figure 1 and the alignment from which the tree was inferred can be found in supplementary table 2.

Taken together, there is no evidence that would support a functional differentiation between cytochromes *c*_6B_ and *c*_6C_. Cytochrome *c*_6B_ is found only in a clade of organisms known for a high protein substitution rate (Dufresne *et al*., 2005), cytochromes *c*_6B_ and *c*_6C_ both have a low redox midpoint potential, share common ancestry to the exclusion of cytochrome *c*_6_, and are not found together in one organism. The distinction between cytochromes *c*_6B_ and *c*_6C_ does not seem to represent functional divergence, and we propose to refer to all as cytochromes *c*_6B_ in future.

### Distribution of cytochrome *c*_6_ family members across photosynthetic eukaryotes

The recent sequencing of genomes and transcriptomes of a wider range of eukaryotic photosynthetic organisms allowed for a more thorough search for *c*_6_-like cytochromes, including cytochrome *c*_6A_. Protein and nucleotide databases across eukaryotes were searched using BLAST for cytochrome *c*_6_ family sequences. Sequences recovered were defined as cytochrome *c*_6A_ if they contained a hydrophobic residue (valine, leucine or isoleucine) at the equivalent of position 52, indicating a low redox midpoint potential, and an insertion containing a disulphide bridge (the LIP) in the loop region compared to cytochrome *c*_6_. Sequences recovered were defined as cytochrome *c*_6B_ if they contained the hydrophobic residue implying a low redox midpoint but not the LIP.

#### Distribution of cytochrome *c*_6_ after primary endosymbiosis

The distribution of cytochromes *c*_6_, *c*_6B_ and *c*_6A_ across cyanobacteria and in eukaryotes after primary endosymbiosis was mapped onto a phylogenetic tree based on an alignment of concatemers of plastid genes and cyanobacterial homologues, figure 5. The presence of cytochromes *c*_6_ and *c*_6B_ in the glaucophyte and red algal lineages suggests that the cyanobacterium involved in the primary endosymbiosis event contained both a cytochrome *c*_6_ and a *c*_6B_.

**Figure 5.**
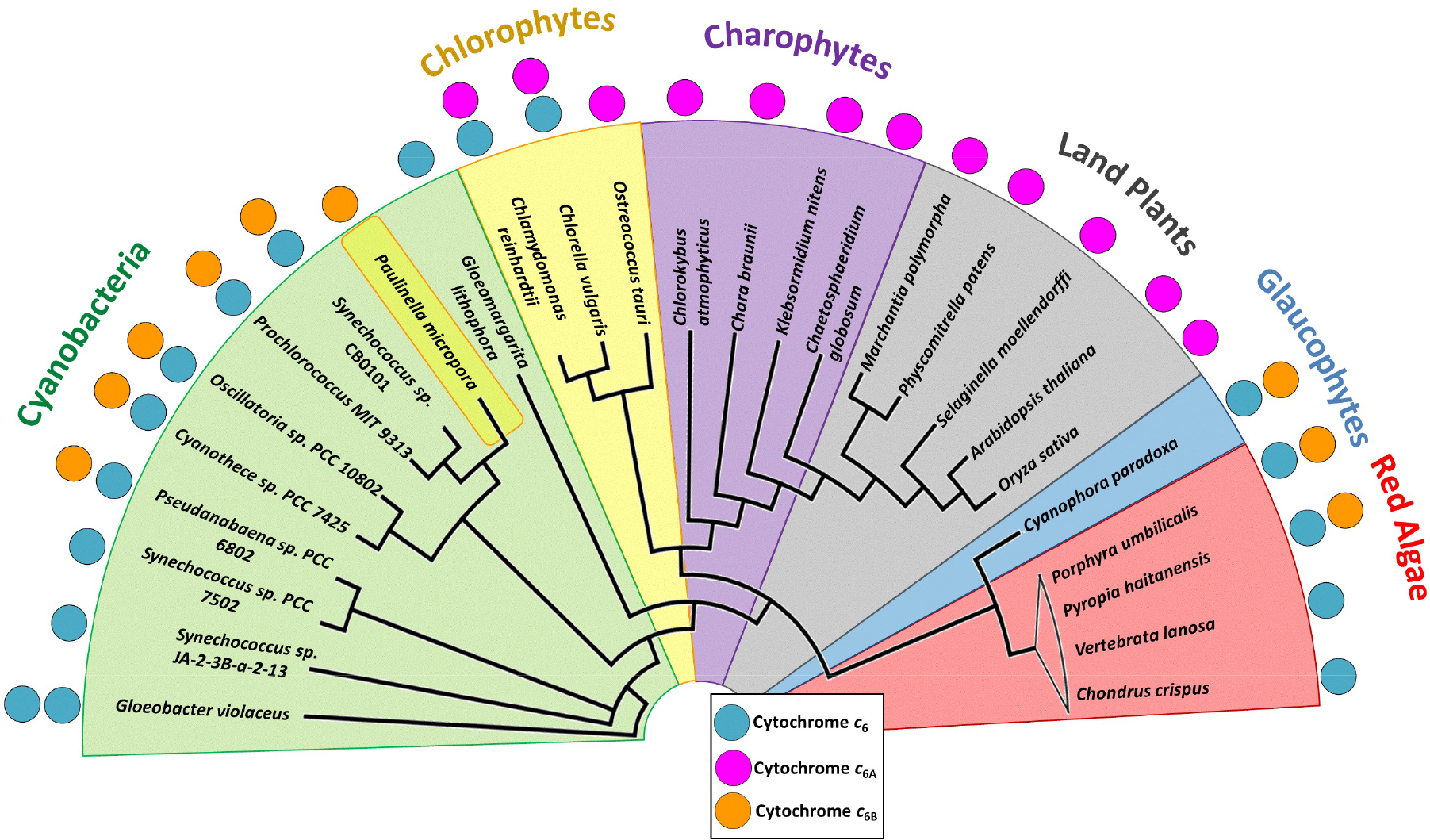
Distribution of *c*_6_-type cytochromes across photosynthetic lineages. Presence of a coloured circle adjacent to a species name indicates that a sequence of the relevant cytochrome was found in sequence database searches, with multiple copies of the same coloured circle indicating a potential paralogue. Paulinella chromatophora is highlighted to segregate it from the cyanobacteria. Phylogenetic tree branch lengths are not to scale. Diagram based on phylogenetic tree from Ponce-Toledo *et al*., 2017. Accession numbers for individual gene sequences can be found in the supplementary table 3.

The results indicate that after primary endosymbiosis, cytochrome *c*_6B_ was replaced by cytochrome *c*_6A_ in the green plant and algal lineage. This suggests that cytochrome *c*_6A_ was derived from cytochrome *c*_6B_, possibly through an insertion of the LIP early in the green chloroplast lineage. The insertion of the LIP into an existing sequence rather than duplication and divergence is supported by the observation that no two species have been found to contain both a cytochrome *c*_6A_ and *c*_6B_ sequence. Although cytochrome *c*_6_ was identified in some chlorophyte species, it was not identified in any charophytes or land plants, suggesting that the loss of cytochrome *c*_6_ occurred in the ancestor to the charophyte lineage. This in turn has resulted in land plants exclusively containing cytochrome *c*_6A_. In contrast, the glaucophytes and many red algal species have retained the original cytochrome *c*_6_.

However, some red algal species (though not all) appeared to have lost cytochrome *c*_6B_, for example in *Chondrus crispus*. Finally the eukaryotic protist *Paulinella* contains a cytochrome *c*_6B_-like sequence. This is likely to reflect the recent, independent primary endosymbiosis event that gave rise to the *Paulinella* chloroplast (Marin, Nowack and Melkonian, 2005; Yoon *et al*., 2006), although the *Paulinella* line also appears to have lost cytochrome *c*_6_.

#### Distribution of cytochrome *c*_6_ family members after secondary endosymbiosis

Many photosynthetic eukaryotes contain chloroplasts of secondary origin. We therefore searched for the presence of cytochromes *c*_6_, *c*_6B_ and *c*_6A_ in these organisms. *Euglena gracilis*, which contains a chloroplast of secondary green origin (Turmel *et al*., 2009), was predicted to contain a cytochrome *c*_6A_ sequence in addition to cytochrome *c*_6_ (Novák Vanclová *et al*., 2020). The chlorarachniophytes, a class of rhizaria with a secondary chloroplast of green origin (Rogers *et al*., 2007), on the other hand only had a cytochrome *c*_6_ sequence and no evidence of cytochrome *c*_6A_.This suggests that either the green algal endosymbiont of the chlorarachniophytes did not have cytochrome *c*_6A_ or that the gene was lost after secondary endosymbiosis.

Different organisms with a secondary red chloroplast also varied in cytochrome *c*_6_ family gene distribution. Haptophytes, cryptomonads and some ochrophytes, which contain a chloroplast of a red algal origin (Yoon *et al*., 2002), contained cytochrome *c*_6_ and *c*_6B_ sequences, as expected. However, many ochrophyta and some haptophytes and cryptomonads had no evidence of *c*_6B_ sequences. This suggests that the red algal endosymbiont to haptophytes, cryptomonads and ochrophytes had retained cytochrome *c*_6B_, and that the gene was lost afterwards downstream in certain lineages, although widespread lateral transfer cannot be excluded.

The situation in dinoflagellate algae is complex. The peridinin dinoflagellates contain a chloroplast of secondary red origin (Dorrell and Howe, 2015). Two peridinin dinoflagellates (*Amphidinium carterae* and *Symbiodinium microadriaticum*) contain a cytochrome *c*_6_ sequence and what appeared to be a cytochrome *c*_6A_ sequence. In contrast, *Karlodinium veneficum*, a fucoxanthin dinoflagellate (which obtained its chloroplast via serial endosymbiosis of a haptophyte (Dorrell and Howe, 2015; Klinger *et al*., 2018)), contains a cytochrome *c*_6_ and cytochrome *c*_6B_. The existence of a cytochrome *c*_6B_ in the *Karlodinium* lineage, whose chloroplast is of red algal origin, is not surprising. However, the peridinin dinoflagellate chloroplast is also of red algal origin, so the apparent existence of cytochrome *c*_6A_ in this lineage is unexpected.

The LIP sequences of sequences resembling cytochrome *c*_6A_ in peridinin dinoflagellate algae were compared to those of other cytochrome *c*_6A_ proteins (figure 6). This revealed that the dinoflagellate LIPs show little sequence similarity with the LIP found in cytochromes *c*_6A_ from the green chloroplast lineage, except for the characteristic conserved cysteine residues. This suggests that the LIP sequences in dinoflagellate cytochromes *c*_6A_ have a functional similarity to other LIPs, but that the dinoflagellates acquired this LIP independently.

**Figure 6.**
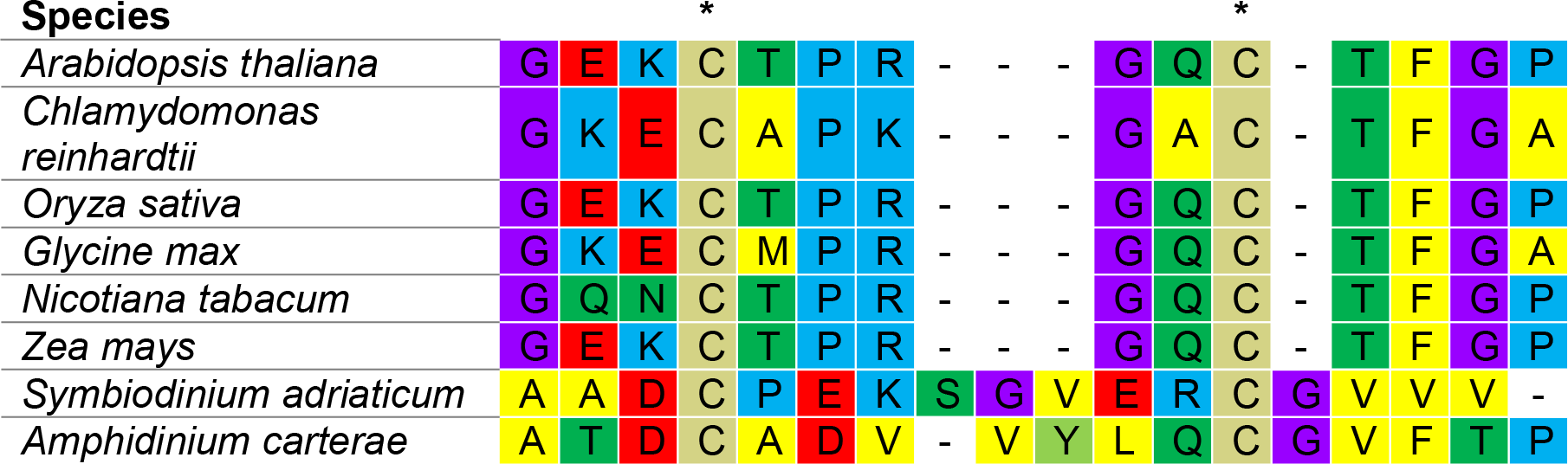
Muscle alignment of loop insertion peptides from cytochromes *c*_6A_ from green eukaryotic lineages and the dinoflagellates *Symbiodinium microadriaticum* and *Amphidinium carterae*. Amino acids are coloured such that yellow-hydrophobic residues, green-polar residues, blue-positively charged residues, beige-cysteines, purple-glycines, light green-tyrosine, and red-negatively charged residues. Dashes indicate inserted gaps. Asterisks represent the two conserved cysteine residues. Accessions used: *A. thaliana* (AT5G45040.1), *C. reinhardti* (XP_001692119.1), *O. sativa* (EAZ04378.1), *G. max* (KRH50430.1), *N. tabacum* (XP_016489567.1), *Z. mays* (ACN28933.1), *S. microadriaticum* (OLP91854.1) and *A. carterae* (CF065358.1).

A summary of the photosynthetic eukaryotes and the presence of each cytochrome *c*_6_ family sequences found is in Table 1, with taxon IDs of searched clades can be found in supplementary table 4 and accession numbers of the found sequences are listed in supplementary table 1.

**Table 1:** The presence or absence of cytochrome *c*_6_ family members across photosynthetic eukaryotes.

| Supergroup | Clade<sup>1</sup> | Plastid | Cytochrome $c_6$<sup>2</sup> | Cytochrome $c_{6A}$<sup>2</sup> | Cytochrome $c_{6B}$<sup>2</sup> |
| --- | --- | --- | --- | --- | --- |
| Glaucophyta | Glaucophyta (n= 1) | Primary red | ✓ (1) | ○ | ✓ (1) |
| Rhodophyta | Rhodophyta (n= 190) | Primary red | ✓ (190) | ○ | ✓ (4) |
| Viridiplantae | Chlorophyta (n= 42) | Primary green | ✓ (32) | ✓ (34) | ○ |
| Viridiplantae | Streptophyta (n= 217) | Primary green | ○ | ✓ (215) | ○ |
| SAR - Rhizaria | *Paulinella* (n= 3) | Primary green | ○ | ○ | ✓ (3) |
|  | Chlorarachniophyta (n= 2) | Secondary green | ✓ (2) | ○ | ○ |
| Cryptomonads | Cryptomonads (n= 7) | Secondary (rhodophyta) | ✓ (2) | ○ | ✓ (1) |
| Haptista | Haptophyta (n= 6) | Secondary (rhodophyta) | ✓ (6) | ○ | ✓ (3) |
| SAR - Heterokonta | Ochrophyta (n= 80) | Secondary (rhodophyta) | ✓ (78) | ○ | ✓ (2) |
| SAR - Alveolata | Dinoflagellata (n= 7) | Secondary (rhodophyta)/ Tertiary (haptophyta) | ✓ (6) | ✓ (2) | ✓ (1) |
| Discoba - Euglenozoa | Euglenids (n= 2) | Secondary (chlorophyta) | ✓ (2) | ✓ (1) | ○ |

<sup>1</sup> Where n represents the number of organisms searched.
<sup>2</sup> The number of sequences found is given in brackets.

### Cytochrome *c*_6A_ arose from cytochrome *c*_6B_ rather than directly from cytochrome *c*_6_

To test if cytochrome *c*_6A_ (non-dinoflagellate) arose from cytochrome *c*_6B_, rather than by independent modification of a cytochrome *c*_6_, a phylogenetic tree was inferred using cytochrome *c*_6A_ sequences from eukaryotic algae and higher plants, together with cytochromes *c*_6_ and *c*_6B_ from a wide range of cyanobacteria (figure 7 shows a condensed version of this tree). Cytochromes *c*_6A_ and *c*_6B_ grouped together to the exclusion of cytochrome *c*_6_ (bootstrap value of 75), suggesting that cytochrome *c*_6A_ shares most recent common ancestry with cytochrome *c*_6B_. This supports the conclusion above (figure 5) that cytochrome *c*_6A_ was derived from cytochrome *c*_6B_ through an insertion event in the loop region. Once again, the bootstrap values in the tree were considerably lower than those of the species tree established by Walter *et al*., 2017, but this is to be expected as the *c*_6_ family cytochrome sequences are short.

**Figure 7.**
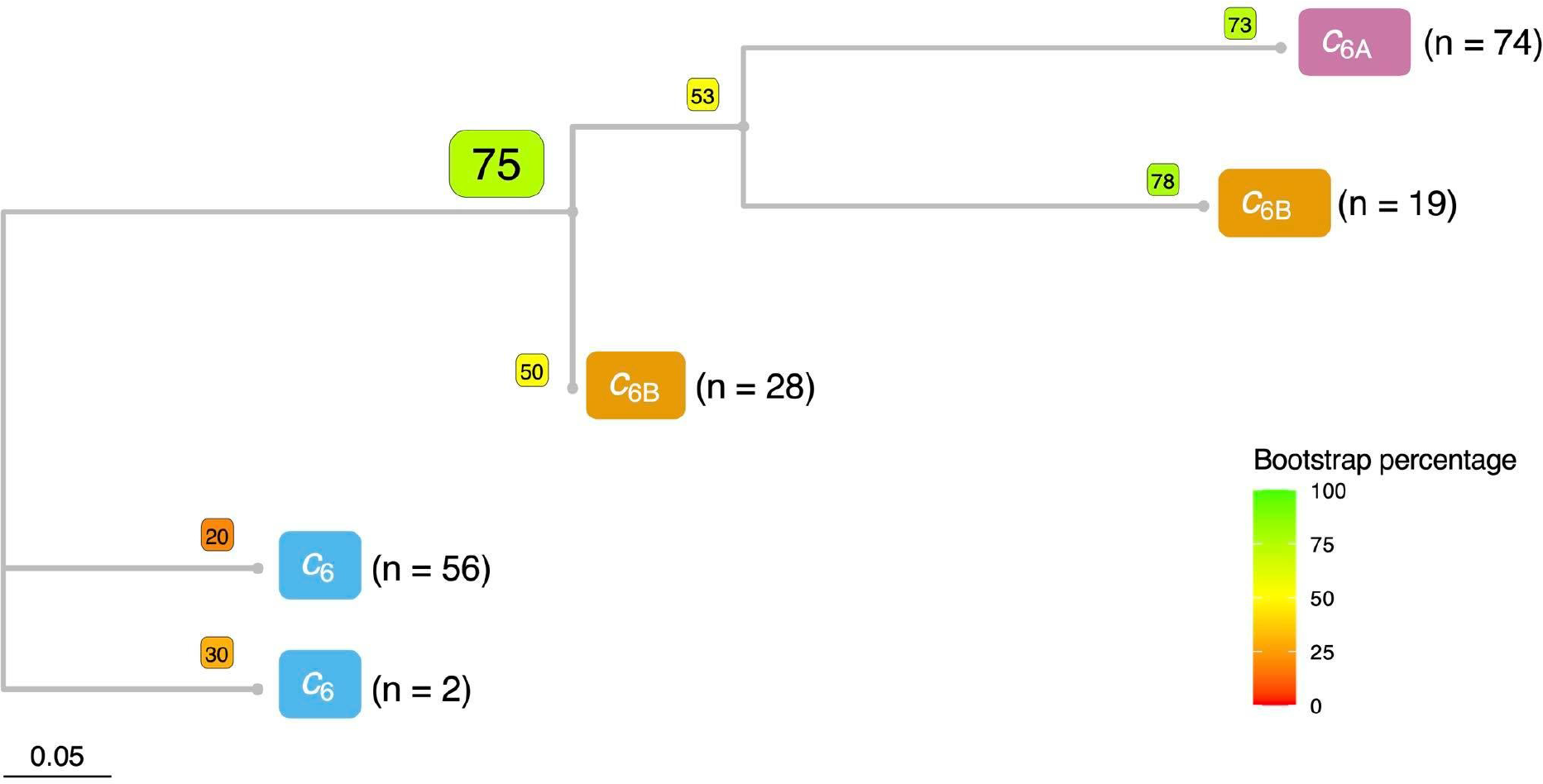
Condensed phylogenetic tree inferred from an alignment of cytochrome *c*_6_, *c*_6A_, and *c*_6B_ peptide sequences from eukaryotic algae, higher plants and cyanobacteria (coloured blue, pink and orange respectively). Alignments were performed using Muscle algorithm and can be found in the supplementary information along with accessions for each sequence used. The tree was built using maximum likelihood inference using a WAG model with Gamma distribution and invariant sites (WAG+G+I). Bootstrap values for each branch point, using 100 iterations, are shown in coloured boxes. The full tree is shown in supplementary figure 2 and the alignment from which the tree was inferred can be found in supplementary table 5.

## Discussion

### Cytochromes *c*_6B_ and *c*_6C_ are orthologues

The original differentiation of cytochromes *c*_6B_ and *c*_6C_ was based on the sequence data available at the time (Bialek *et al*., 2008). However, now that more genomic sequence data are available, cytochrome *c*_6_ family sequences from a larger range of taxa can be analysed. Our analysis indicates that the distinction between cytochromes *c*_6B_ and *c*_6C_ can be accounted for by taxon sampling rather than differences in function. (Although cytochromes *c*_6B_ and *c*_6C_ lie on opposite sides of the root of the cytochrome *c*_6_ family in the tree of Bialek *et al*. (2008), the placing of the root should be viewed with caution given that it depends on *c*-type cytochromes of very different function from the cytochrome *c*_6_ family.) In addition, as the crystal structures, surface charge distribution and redox midpoint potentials of cytochromes *c*_6B_ and *c*_6C_ are notably similar (Bialek and Jaskolski, 2014; Zatwarnicki *et al*., 2014b; figure 1), it seems likely that cytochromes *c*_6B_ and *c*_6C_ perform a similar function and are thus orthologues.

### Two independent origins of *c*_6A_

Although the presence of cytochrome *c*_6A_ in plants and green algae has been known for time, the presence of cytochrome *c*_6A_ in peridinin dinoflagellates was unexpected. Dinoflagellates contain chloroplasts of secondary or tertiary origin, depending on species. The chloroplast found in *Symbiodinium microadriaticum* and *Amphidinium carterae* contains peridinin, and is believed to represent the ancestral dinoflagellate chloroplast. This chloroplast was most likely obtained through secondary endosymbiosis of red algae (Dorrell and Howe, 2015). Therefore, these species might be expected to contain a cytochrome *c*_6B_. Instead, the peridinin dinoflagellates contain a cytochrome *c*_6A_-like sequence. Two hypotheses for this are a) the result of lateral gene transfer from an organism with cytochrome *c*_6A_ and the loss of the cytochrome *c*_6B_ or b) the insertion of a LIP-like sequence into an existing cytochrome *c*_6B_ sequence. Although lateral gene transfer to dinoflagellates from other organisms has been well documented (Takishita, Ishida and Maruyama, 2003; Hackett *et al*., 2005; Chan *et al*., 2012; Wisecaver, Brosnahan and Hackett, 2013), the low sequence similarity between the dinoflagellate *c*_6A_ LIP and those from the green plant lineage would suggest an independent LIP insertion into cytochrome *c*_6B_ is more likely.

### The current model of cytochrome *c*_6_ family ancestry

The analysis of the cytochromes *c*_6_, *c*_6A_ and *c*_6B_ in this study has provided an updated model of the protein family’s ancestry (figure 8). As more anciently diverged cyanobacterial species such as *Gloeobacter* appear to contain cytochrome *c*_6_ exclusively, this suggests that the low redox midpoint potential cytochromes are more recent than cytochrome *c*_6_. A duplication of cytochrome *c*_6_, followed by point mutations that lowered the redox midpoint potential, led to the evolution of cytochrome *c*_6B_. This is supported by the presence of both cytochromes *c*_6_ and *c*_6B_ in most extant cyanobacteria today. At primary endosymbiosis, giving rise to the red, green and glaucophyte chloroplasts, the genes were transferred to photosynthetic eukaryotes.

**Figure 8.**
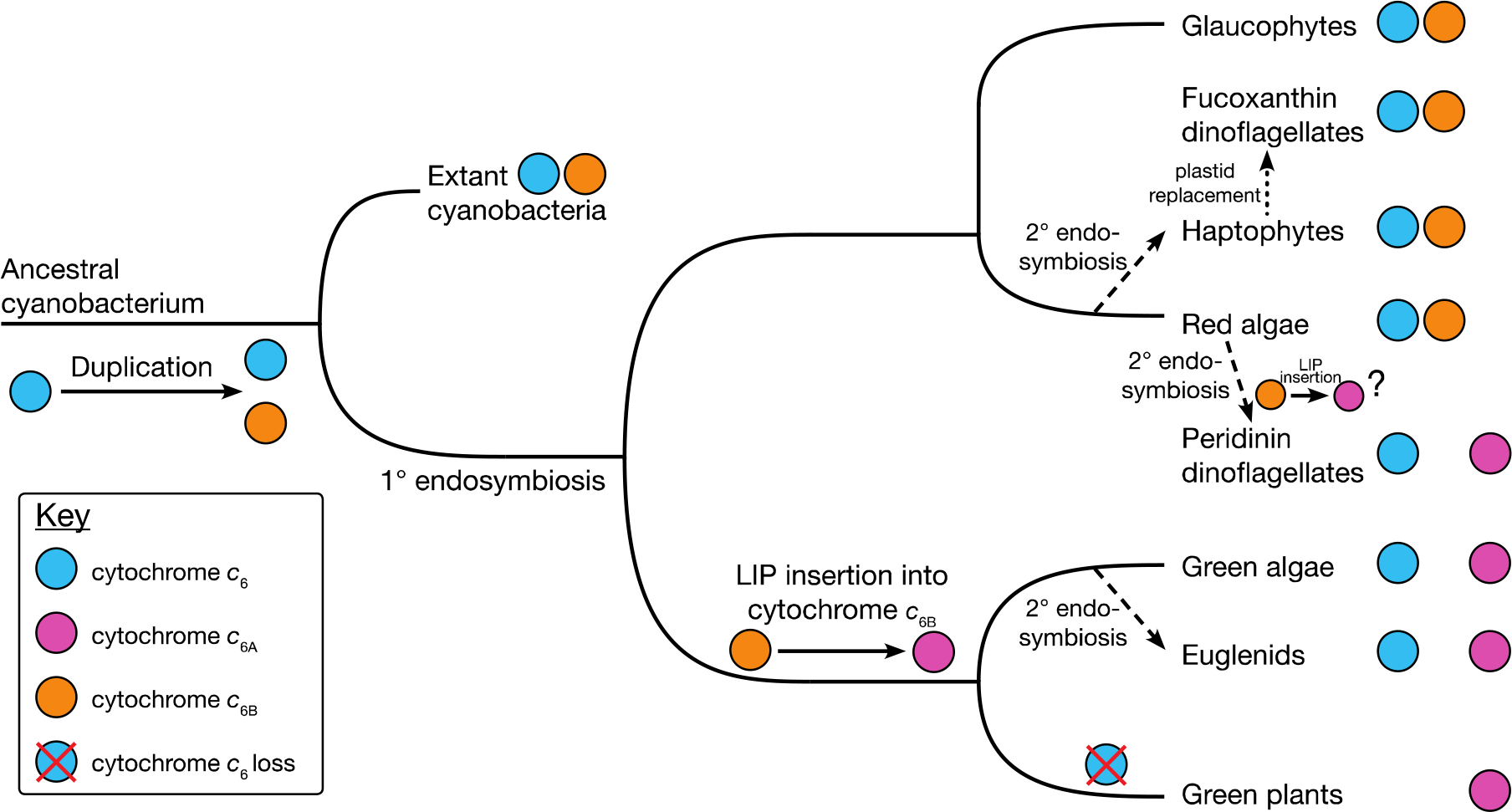
Updated ancestry model for the cytochrome *c*_6_ family in photosynthetic organisms.

Red algal lineages and the glaucophytes contain cytochromes *c*_6_ and *c*_6B,_ although some red algal species have lost cytochrome *c*_6B_. In the green chloroplast lineages, cytochrome *c*_6B_ was replaced by cytochrome *c*_6A_. This was probably due to an insertion of the LIP into cytochrome *c*_6B_, as cytochrome *c*_6A_ is monophyletic within cytochrome *c*_6B_ (figure 7). In many chlorophyte species, both cytochromes *c*_6_ and *c*_6A_ are present. In the charophytes, ancestors to the green land plants, cytochrome *c*_6_ was lost. In consequence, land plants contain only a cytochrome *c*_6A_.

Organisms containing chloroplasts of secondary origin appear have inherited their cytochrome *c*_6_ family genes from the relevant endosymbiont. The haptophytes obtained both cytochromes *c*_6_ and *c*_6B_ from the red algal chloroplast, and these genes were transferred to the fucoxanthin dinoflagellates following serial endosymbiosis. In contrast, the peridinin dinoflagellates, containing chloroplasts of secondary red origin, converted the cytochrome *c*_6B_ into a cytochrome *c*_6A_-like protein through the insertion of a novel LIP.

With green plastid secondary endosymbiosis, genes for both cytochromes *c*_6_ and *c*_6A_ were passed to the euglenids (Novák Vanclová *et al*., 2020).

Overall, it is clear that the low potential cytochrome *c*_6AB_ family is widely, but not universally, present among oxygenic photosynthetic organisms. It is unlikely to be essential under all conditions, but there is no obvious environmental feature common to those organisms that retain a member of the family. The function of the cytochrome *c*_6AB_ family remains to be determined.

## Materials and methods

### Construction of phylogenetic trees

A cytochrome *c*_6B_ sequence (BAD79758.1) was used for searching the “non-redundant protein sequences (*nr*)” database with the NCBI BLASTp algorithm (https://www.ncbi.nlm.nih.gov/), limiting the results to the organisms used in two independent phylogenetic analyses of the cyanobacterial lineage (Schirrmeister *et al*., 2015; Walter *et al*., 2017). A cytochrome *c*_6A_ protein sequence (AED95193.1) was used to search the *nr* protein database and the “nucleotide collection (*nr/nt*)” nucleic acid database with the BLASTp and tBLASTn algorithms respectively, limiting the results to the orders used in a phylogenetic analysis of green plants (Ruhfel *et al*., 2014). The resulting sequences of both searches were downloaded from the NCBI BLAST result page.

The retrieved peptide sequences were imported into MEGA7 (Kumar *et al*., 2016). Sequences were deleted from the selection if they were too short or too long to be a valid cytochrome *c*_6(A/B)_ sequence (less than 80 and more than 200 amino acids before N-terminal targeting peptide trimming) or did not have a CxxCH haem binding motif. The sequences were aligned using the MUSCLE algorithm with UPGMA clustering method and a gap opening penalty of −2.9 and no gap extension penalty. Subsequently the putative signal peptides were trimmed from the sequences.

The WAG model with gamma distribution and invariant sites (WAG+G+I) (Whelan and Goldman, 2001) was determined to be optimal for tree inference with Maximum Likelihood (ML) by the “Find Best DNA/Protein Models (ML)” tool in MEGA7 and was thus used in the algorithm parameters for tree inference. Statistical testing was performed using the bootstrap method with 100 iterations. The final trees were visualised using Treeio in Rstudio (Wang *et al*., 2019). The accessions of the sequences used for inference of phylogenetic trees can be found in supplementary tables 2 and 5.

### Database queries for peptide sequences

Searches for protein sequences homologous to the cytochrome *c*_6_ family, or nucleotide sequences encoding them, were performed using NCBI BLAST both in BLASTp and tBLASTn (https://www.ncbi.nlm.nih.gov/). The cytochrome *c*_6_ *c*_6A_, *c*_6B_ and *c*_6C_ peptide sequences (without targeting) used in both BLASTp and tBLASTn searches were from accessions ALJ67080.1, AED95193.1, AAP99622.1 and ACB00369.1 respectively. For BLASTp searches the database searched was “Non-redundant protein sequences (nr)” with default parameters. For tBLASTn searches the databases searched were “nucleotide collection (nr/nt)”, “Whole-genome shotgun contigs (wgs)” and “Expressed sequence tags (est)” with default parameters. Having identified a putative cytochrome *c*_6_ sequence, each organism that provided a query sequence was searched again to confirm the query as the best hit.

## Supporting information

Supplementary figure 1 alignment

Supplementary figure 2 alignment

## Acknowledgements

This work was supported by a Biotechnology and Biological Sciences Research Council DTP PhD studentship (BB/M011194/1 to B.S.); the German Academic Scholarship Foundation to D.K.; the Gates Cambridge Trust to D.K.; the Benn W Levy Trust to D. K. and the Gordon and Betty Moore Foundation (GBMF4976 doi: https://doi.org/10.37807/GBMF4976 and GBMF9358 doi: https://doi.org/10.37807/GBMF9358 to R.E.R.N. and C.J.H.).

B.S. and D.K. contributed equally to the content of this article.

## Data availability

The data underlying this article are available at the NCBI. References for individual genes are given in the article and/or in the supplementary material. All alignments are provided in the online supplementary material.

