## Supplementary figure 1 legend for "The evolution of the cytochrome *c*_6_ family of photosynthetic electron transfer proteins"

Supplementary figure 1: Phylogenetic tree inferred from an alignment of cytochrome $c_6$, $c_{6B}$ and $c_{6C}$ peptide sequences from cyanobacterial species (coloured blue, orange and green respectively). Alignments were performed using Muscle algorithm. The tree was built using maximum likelihood inference using a WAG model with Gamma distribution and invariant sites (WAG+G+I). Bootstrap values for each branch point, using 100 iterations, are shown in coloured boxes. The alignment from which the tree was inferred can be found in supplementary table 2 and the condensed version of this tree is shown in figure 4.
