## Supplementary figures and images for "The evolution of the cytochrome *c*_6_ family of photosynthetic electron transfer proteins"

### Supplementary figure 1

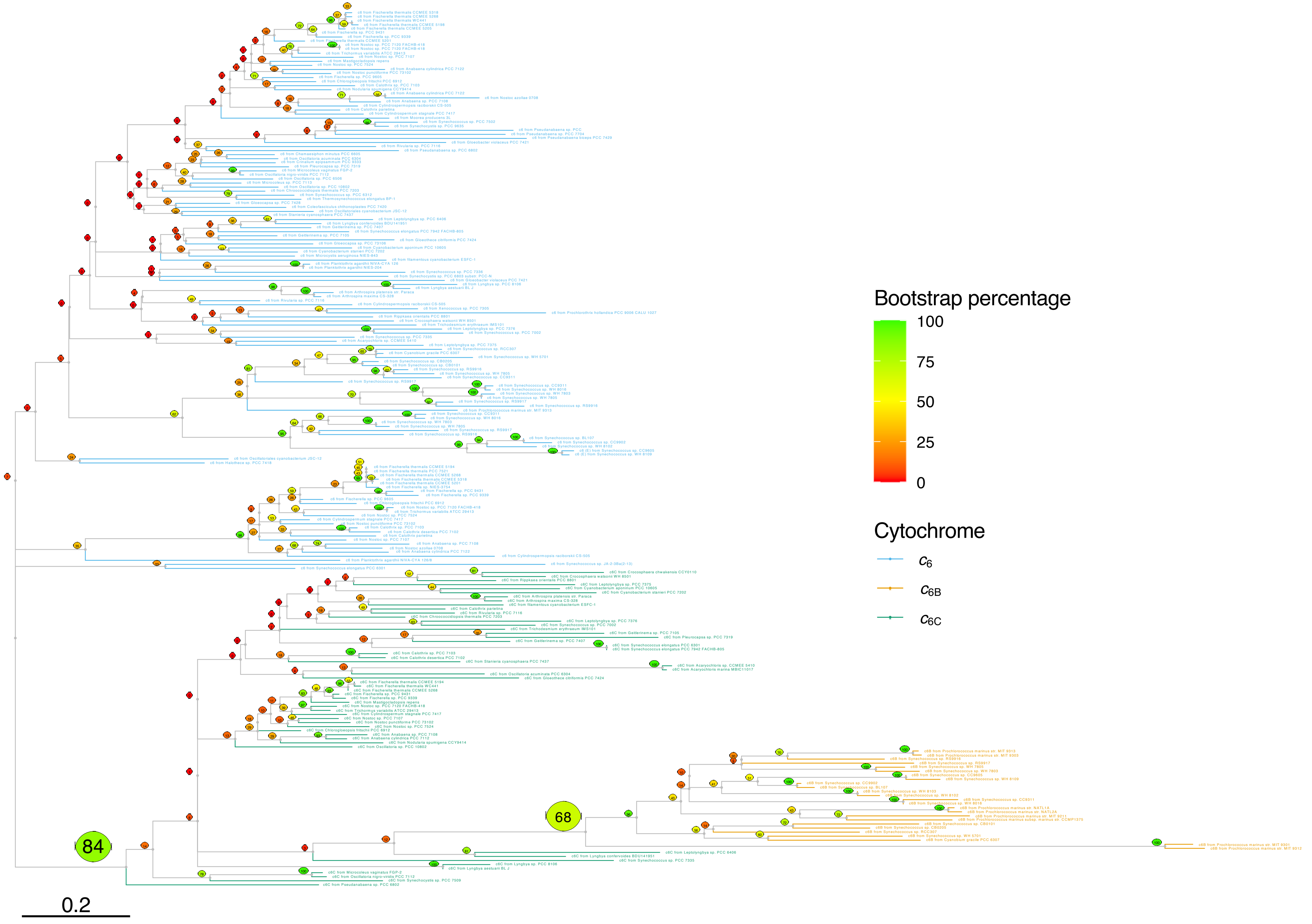

### Supplementary figure 2

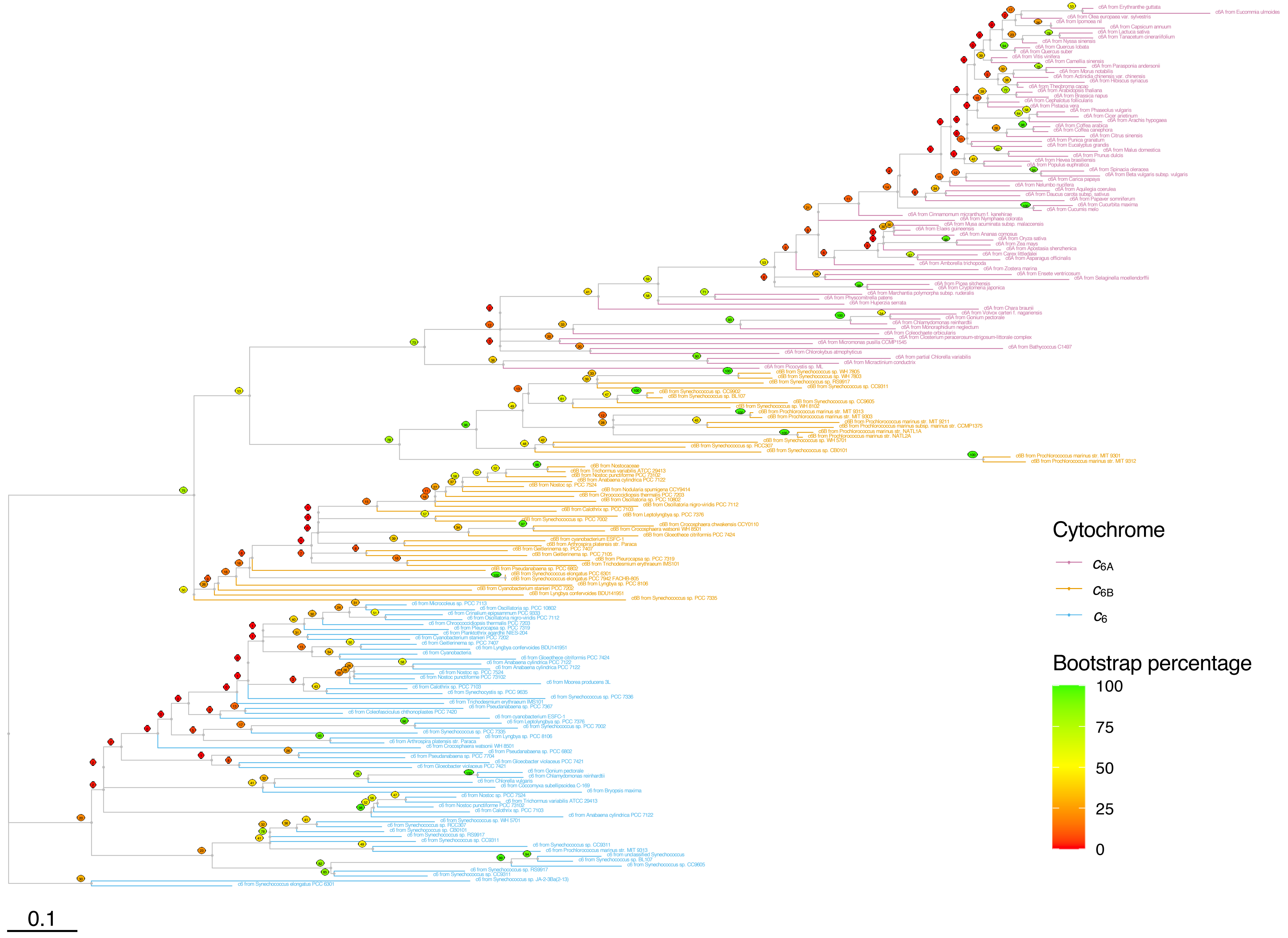
